# Blockade of Tim-3 pathway in a mouse model of Toxoplasmosis: impact on brain leukocyte infiltration, parasite burden, and neuroinflammation

**DOI:** 10.64898/2026.04.06.716688

**Authors:** Raphael P. Viscidi, Jing Huang, Ye Li, Emily G. Severance, Jianchun Xiao

## Abstract

Cell-mediated immune responses are crucial for protecting the host against *Toxoplasma gondii* infection. However, impaired immunity, such as T-cell exhaustion, is a common phenomenon during chronic infection. This may represent a strategy employed by *T. gondii* to evade host defenses. T-cell immunoglobulin and mucin-domain containing 3 (Tim-3) is an important regulatory molecule involved in cell-mediated immunity. This study examined the expression of Tim-3 and the effects of its blockade in a mouse model of toxoplasmosis. In mice with chronic *T. gondii* infection, we found that Tim-3 is highly expressed in both cyst-bearing and non-cyst-bearing tissues, and its expression correlates with the parasite burden. Blocking the Tim-3 pathway with an anti-Tim-3 antibody enhances the immune response, resulting in elevated levels of cytokines (IFN-γ, IL-12p70, IL-2, IL-9) and the chemokine CXCL1 in the serum, increased leukocyte infiltration (CD3+, CD14+ cells) in the brain, and downregulation of Tim-3 expression in microglial cells. As a result, the anti-Tim-3 treatment resulted in a 62% reduction in the number of tissue cysts and a trend towards an increase in the homeostatic signature, P2RY12, in microglia. Our study provides proof of concept for an anti-Tim-3 approach in treating chronic *T. gondii* infection and potentially other brain-residing pathogens.

## Introduction

Chronic infection with *Toxoplasma gondii*, a common neurotropic parasite, poses a constant and severe danger in situations of immunodeficiency or when the parasite is congenitally acquired ^1–4^. Clinical manifestations may include encephalitis, pneumonitis, chorioretinitis, meningitis, and widespread organ involvement in immunocompromised individuals. Individuals with congenital toxoplasmosis are likely to experience neurological and ocular issues, as well as developmental disabilities. Additionally, latent toxoplasmosis has been epidemiologically associated with various neuropsychiatric disorders, including autism, cognitive deficits, schizophrenia, and Alzheimer’s disease ^5–8^. Available anti-*T. gondii* drugs, such as sulfadiazine, spiramycin, and pyrimethamine, can effectively suppress active infections caused by tachyzoites. However, they fail to eliminate chronic infections characterized by tissue cysts ^9^. Infection with these cysts remain incurable to date, highlighting the need for innovative therapies.

Cell-mediated immune responses by resident central nervous system (CNS) and infiltrating peripheral immune cells are essential for protection against chronic *T. gondii* infection, especially in preventing tissue cyst reactivation and toxoplasmic encephalitis ^10^. Interferon-gamma (IFN-γ) is a key mediator of this protection, produced by CD8+ and CD4+ T cells. Microglia activation is characteristic of chronic *T. gondii* infection, as these cells are resident immune cells of the CNS ^11,12^. Microglia have various roles in maintaining homeostasis and promoting inflammation in numerous disease conditions ^13^. Microglia lose their usual homeostatic transcriptional profile under pathological circumstances, such as neuroinflammation. The purinergic receptor P2RY12 is commonly recognized as a marker of non-activated, homeostatic microglial cells ^14^. P2RY12 expression is high in the “resting” state but decreases significantly following microglial activation ^15^. For example, P2RY12 levels decline after events such as brain injury, aging, multiple sclerosis, and Alzheimer’s disease, where microglia shift to an activated phenotype ^13^.

Previous studies indicate that T cell exhaustion is common in chronic *T. gondii* infection, significantly contributing to disease progression ^16,17^. This exhausted state results from prolonged antigen exposure and has been characterized by reduced proinflammatory cytokine production and decreased cytotoxicity ^18^. Exhausted T cells also exhibit sustained expression of inhibitory receptors that deliver negative signals to prevent T cell activation. In mouse models of *T. gondii* infection, several inhibitory receptors—such as PD-1, CTLA-4, TIGIT, Tim-3, LAG-3, and 2B4—are expressed by CD4+ and CD8+ T cells ^19^. Bhadra et al. showed that in genetically susceptible C57BL/6 mice, blocking the PD-1 ligand (PD-L1) can reactivate exhausted CD8 T cells and prevent parasite recrudescence ^17^. Previous research from our laboratory has demonstrated that inhibiting the PD-1–PD-L1 interaction in genetically resistant mouse models can reverse antigen-specific CD8 T cell exhaustion, eliminate tissue cysts, and reduce neuroinflammation ^20,21^.

T-cell immunoglobulin and mucin-domain containing protein 3 (Tim-3) is expressed in various immune cell types, including IFN-γ-producing CD4+ and CD8+ T cells, regulatory T cells (Treg cells), myeloid cells, natural killer cells, and mast cells ^22^. Tim-3 plays a crucial role in regulating both adaptive and innate immunity. In adaptive immunity, CD4+ T cells can differentiate into Th1 and Th2 cells depending on the type of pathogens encountered. Tim-3 has been identified as a Th1-specific marker and negatively regulates the Th1 response by binding to its ligand, galectin 9. This process is commonly involved in multiple diseases, including autoimmunity, infection, and cancer ^23,24^. Tim-3 expression indicates a highly dysfunctional or terminally exhausted subset of CD8+ T cells in cancer ^25,26^, as well as in patients infected with viruses such as HIV, HCV, HBV, and LCMV ^24^. Blocking Tim-3 with antibodies has been shown to restore T cell proliferation, enhance cytokine production, and provide benefits in experimental models of various infections and T cells from chronically infected patients ^27,28^.

Host resistance to *T. gondii* relies on Th1-mediated immunity ^10^, highlighting the importance of CD4 T cells. In AIDS patients, reactivation of *T. gondii* infection often occurs when CD4 T cell counts drop below 100 cells/mm³ ^29^. Moreover, studies have shown that CD4 T cell exhaustion leads to the dysfunction of CD8 T cells ^30^. The potential to block the Th1-specific marker Tim-3 in chronic *T. gondii* infection is unexplored, though increased expression has been noted in the brain and spleen ^31,32^. We hypothesized that blocking Tim-3 would affect the level of chronic *T. gondii* infection and the immune response against the parasite. Specifically, we examined parasite clearance and immune markers in both the periphery and the brain following Tim-3 blockade.

## Methods

### Mouse model of chronic T. gondii infection

The protocol was approved by the Animal Care and Use Committee at Johns Hopkins University. All experiments conformed to the U.S. National Institutes of Health Guide for the Care and Use of Laboratory Animals, and all methods are reported in accordance with ARRIVE guidelines. We used a previously established chronic mouse model of infection ^33^. The model employs a type I strain. We chose this strain because of its close association with clinical disease and the influence on host genes demonstrated in our previous research ^34–37^. In brief, 8-week-old female CD-1 mice (ICR-Harlan Sprague) were infected intraperitoneally (i.p.) with 500 tachyzoites of the *T. gondii* GT1 strain. To control tachyzoite proliferation and prevent mortality during the acute phase, the mice were treated with sulfadiazine sodium in drinking water (400 mg/L) from days 5 to 30 post-infection. Control mice received only the vehicle (PBS). Tim-3 blockade was initiated at 14 weeks post-infection (wpi) as described below. Tissue samples from the prefrontal cortex, striatum, heart, liver, and spleen were collected to analyze Tim-3 expression as part of a previously published study ^33^.

### Infection characterization

Infection was characterized serologically by the presence of IgG antibodies against the entire *T. gondii* organism as well as peptide antigens from the *T. gondii* cyst protein, MAG1. The anti-*T. gondii* antibodies were measured using a modified commercial ELISA kit, while the anti-MAG1 antibodies were determined using a previously developed MAG1 ELISA assay ^38,39^. The serum was diluted at 1:100 for antibody testing.

### In Vivo Blockade of the Tim-3 Pathway

To block the Tim-3 pathway, 100 μg of rat anti-mouse Tim-3 antibody (αTim-3, RMT3-23, BioLegend) was administered intraperitoneally every 2 days for 2 weeks. The effectiveness of αTim-3 in blocking the Tim-3 pathway has been demonstrated previously ^40^. An isotype control of 200 μg rat anti-mouse IgG2b (BioLegend) was administered as described in a previous study^20^. The use of a higher dose of isotype control is to provide a more robust negative control, which ensures that non-specific Fc-mediated effects are fully accounted for ^40,41^. Mice were sacrificed 2 weeks after the final injection, and blood samples and brain tissue were collected.

### Serum cytokine quantification

At the time the mice were sacrificed, blood samples were collected, and serum was isolated. Levels of 23 cytokines and chemokines in the serum were measured using Bio-Plex multiplex assay (Bio-Rad). Analytes measured included Eotaxin, G-CSF, GM-CSF, IFN-γ, IL-1α, IL-1β, IL-2, IL-3, IL-4, IL-5, IL-6, IL-9, IL-10, IL-12p40, IL-12p70, IL-13, IL-17A, KC (CXCL1), MCP-1, MIP-1α, MIP-1β, RANTES, and TNF-α. All samples were tested following the manufacturer’s instructions.

### Cyst enumeration

The brain was cut sagittally along the midline. One half of the brain tissue was used to prepare homogenates, while the other half was used for immunohistochemical analyses. Tissue cysts in the brain homogenates were enumerated as described previously ^33^. In brief, samples were examined using a fluorescence microscope in seven 8-μL samples. The reported numbers reflect the counts from the half-brain multiplied by two. The size of each cyst was determined by measuring its maximum diameter.

### Quantitative PCR

RNA was extracted from mouse tissues and brain homogenates. Quantitative PCR was performed for Tim-3 (TaqMan, Life Technologies), BAG1 ^42^, and SAG1 ^43^ transcripts. A sample was considered positive if a signal was detected at < 40 cycles. For Tim-3, the fold changes between groups were assessed using relative quantification (delta-delta Ct method), with β-actin serving as the endogenous control. For BAG1, qPCR was performed three times for each mouse, and results were scored as positive or negative. The percentage of times the test was positive was calculated for each mouse.

### Immunoblot analysis

Total protein was extracted from brain homogenates using T-PER™ Tissue Protein Extraction Reagent (Thermo Scientific) containing protease inhibitors. A volume of 10∼40 μg of total protein was loaded onto a 4–20% TGX protein gel (Bio-Rad) for electrophoresis under reducing conditions and then transferred to PVDF membranes (Bio-Rad). The membranes were blocked with Starting Block T20 (TBS) Blocking Buffer (Thermo Scientific) for 1 h at room temperature (RT), followed by incubation with primary antibodies at 4 °C overnight. Proteins were probed with primary antibodies for CD3 (monoclonal, cat. 14-0032-85, 1:500, Thermo Scientific), CD14 (polyclonal, cat. 17000-1-AP, 1:500, Thermo Scientific), P2RY12 (monoclonal, cat. 848002, 1:400, Biolegend), and IBA1 (polyclonal, cat. 016-20001, 1:400, Wako). The CD-3 probe blots were stripped and re-probed with anti-CD14 antibodies, while the P2RY12 probe blots were stripped and re-probed with anti-IBA1 antibodies. Bands were visualized using enhanced chemiluminescence (SuperSignal West Femto Maximum Sensitivity Substrate, Thermo Scientific). Protein values were normalized for corresponding values of β-actin. Relative optical density was determined using ImageLab software (Bio-Rad).

### Immunofluorescence staining

Paraffin-embedded tissue sections were cut into a thickness of 5 μm. The sections were treated with xylene and a graduated series of alcohol and rinsed in distilled water. Antigen retrieval was performed using VisUCyte Antigen Retrieval Reagent-Basic (VCTS021, R&D Systems) at 92–95 °C for 20 minutes. After blocking with BlockAid buffer (B10710, Fisher Scientific) for 30 min at RT, the sections were stained overnight with the following primary antibodies: anti-IBA1 (polyclonal, cat. 019-19741, 1:200, Wako), anti-Tim-3 (monoclonal, cat. MCA5790GA, 1:100, Bio-Rad), anti-P2RY12 (monoclonal, cat. 848002, 1:100, Biolegend), and anti-Tim-3 (monoclonal, cat. ab241332, 1:200, Abcam). Secondary antibodies were purchased from Thermo Scientific. Finally, the sections were stained with ProLong™ Diamond Antifade Mountant with DAPI (Thermo Scientific). Three mice were used per group, with 5 sections each. Images were visualized using an Olympus BX41 upright microscope, equipped with cellSens Acquisition software and an Imaging RET IGA6000 monochrome camera. The mean fluorescence intensity (MFI) and colocalization (plugin JACoP) were determined from eight representative images using ImageJ ^44^. Blood cell staining were removed prior to analysis, and the threshold was manually set following visual inspection and remained constant for the same analysis combinations.

### Immunohistochemistry

For immunohistochemistry staining, antibodies against CD3 (monoclonal, cat. 81324-1-RR, 1:1000, Proteintech), CD14 (Polyclonal, cat. 17000-1-AP, 1:1000, Proteintech), and IHC Prep & Detection Kit for Rabbit Primary Antibody (cat. PK10017, Proteintech) were used. Following deparaffination with xylene and rehydration, antigen retrieval was performed by boiling at 95–98 °C for 15 minutes. Slides were incubated overnight at 4 °C with the primary antibodies in a blocking buffer. Staining was revealed by the DAB substrate kit (cat. ab64238, Abcam). A negative control was included by omitting the primary antibody, while the positive control used was spleen tissue.

### Statistical Analysis

Data are presented as means ± SD unless otherwise specified. All data sets were assessed for normality using the Shapiro-Wilk test and for variance using the F-test. Differences between groups were analyzed using Student’s t-test if the data are normally distributed, or the Mann-Whitney test if the data are not normally distributed. For multiple groups, ANOVA with Bonferroni’s correction for multiple comparisons was applied. For monitoring weight change, a repeated-measures ANOVA was applied with the group as the between-subjects factor and time as the within-subjects factor. Because immune checkpoint inhibitor blockade is expected to activate the immune system and increase cytokine production ^45,46^, the one-tailed test was applied for cytokine analyses. To perform statistical analysis, cytokine levels not detectable by the Bio-Plex assay were assigned a value of half the lowest standard concentration. Correlation analysis was performed using two-tailed Spearman’s correlation coefficient (r). Significance was denoted as p < 0.05. Statistical analyses were conducted in Graph-Pad Prism V10.4.2.

## Results

### Tissue cyst-associated elevation of Tim-3 expressions in multiple tissues

As previously described ^33^, the model of outbred CD-1 mice infected with the virulent *T. gondii* strain generated 4 distinct phenotypes based on *T. gondii* IgG and MAG1 antibody profiles, reflecting a range of infection severity. These phenotypes are categorized as follows: (1) IgG+/MAG1+ low (MAG1 < 0.5), representing 24% of the infected mice; (2) IgG+/MAG1+ high (MAG1 ≥ 0.5), 22% of the infected mice; (3) IgG+/MAG1-, 26% of the infected mice; and (4) IgG−/MAG1−, 28% of the infected mice. *T. gondii* IgG is a marker of exposure to the parasite, while MAG1 is a marker for cyst burden ^33^.

We analyzed the mRNA expression of Tim-3 in various mouse tissues, including those with and without cysts. Among the 4 groups, only mice with high levels of MAG1 antibodies showed a significant increase in Tim-3 expression compared to the control group (Fig. 1A). Specifically, Tim-3 mRNA levels increased by 3.1-fold in the prefrontal cortex, 3.9-fold in the striatum, 3.9-fold in the heart, and 2.3-fold in the liver when compared to the control group. There was no significant change in the spleen. Additionally, MAG1 antibody levels were positively correlated with the upregulation of Tim-3 in the prefrontal cortex (r = 0.72, p = 0.0015), striatum (r = 0.81, p = 0.0002), heart (r = 0.59, p = 0.0152), and liver (r = 0.61, p = 0.0113). Conversely, a negative correlation was observed in the spleen (r = -0.51, p = 0.04) (Fig. 1B).

**Figure 1.**
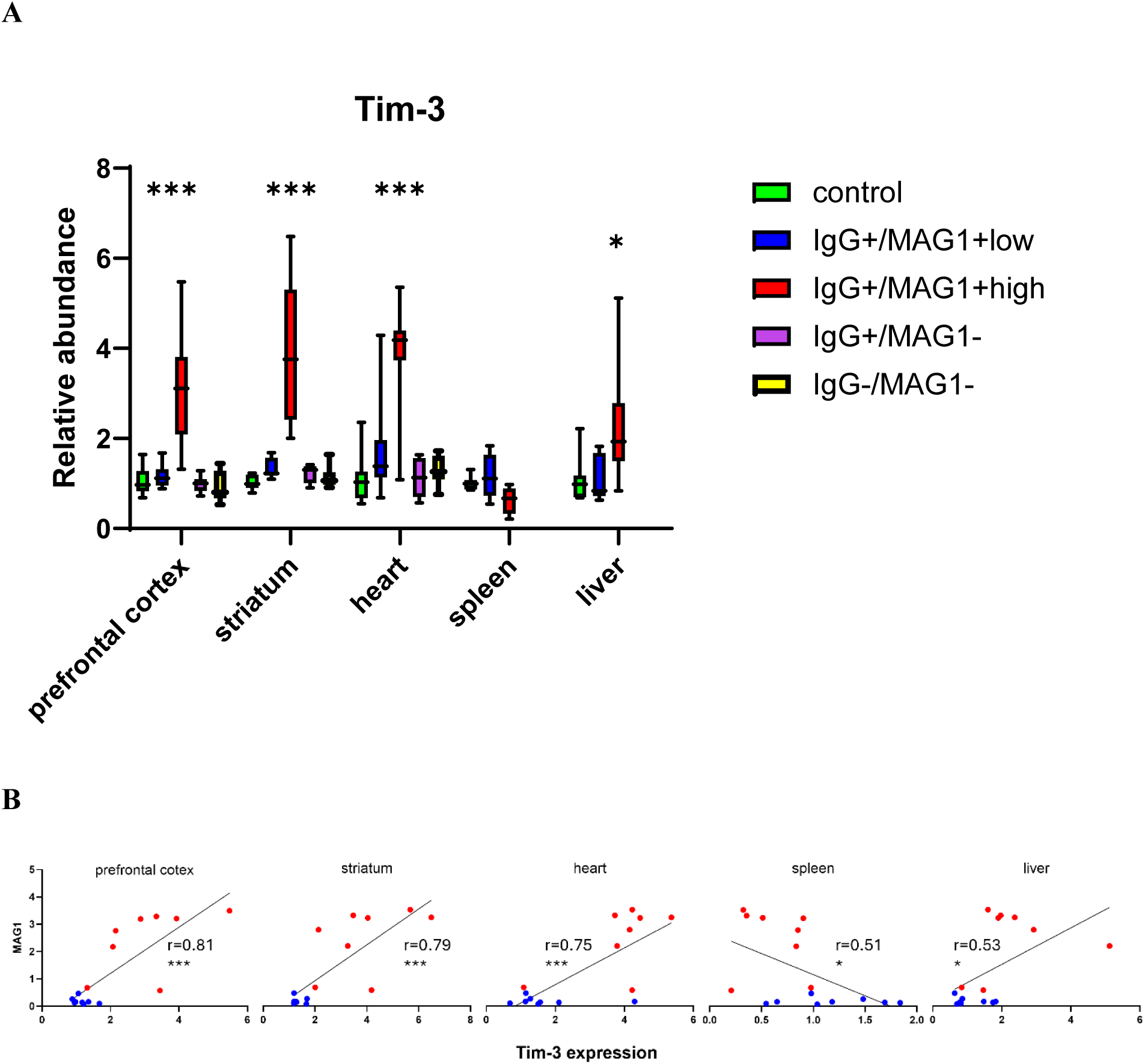
mRNA expression of Tim-3 in multiple tissues of mice exposed to *T. gondii* and its correlation with MAG1 antibody levels. **(A)** Levels of Tim-3 mRNA expression in multiple tissues from mice with different antibody profiles measured by RT-qPCR (n=8 per group). The box and whisker plot illustrates the distribution of delta Ct values: the bottom and top of the box represent the first and third quartiles, respectively, while the line inside the box indicates the median. The ends of the whiskers represent the minimum and maximum values in the data. IgG refers to antibodies against *T. gondii*, while MAG1 refers to antibodies against the *T. gondii* MAG1 antigen. Presence is indicated by “+” and absence by “−”. MAG1 antibody levels further divided into high (OD > 0.5) and low (OD < 0.5) groups. Statistical analysis was conducted using ANOVA followed by Bonferroni’s post hoc test. **(B)** Spearman’s correlation analysis between Tim-3 mRNA expression levels and the MAG1 antibody levels in both IgG+/MAG1+high and IgG+/MAG1+low groups. *p < 0.05, ***p < 0.001. Red dot: MAG1-high; blue dot: MAG1-low.

### Blockade of Tim-3 reduced the number of brain tissue cysts

Since Tim-3 was found to be overexpressed only in mice with high levels of the MAG1 antibody, we selected MAG1-high mice for the treatment with anti-Tim-3 antibody. The mice were randomly divided into two groups: one receiving anti-Tim-3 (αTim-3) and the other receiving isotype control, with 5 mice in each group. The MAG1 antibody levels were similar between the two groups before treatment (αTim-3: MAG1 = 3.02 ± 0.61; isotype: MAG1 = 2.87 ± 0.54; p = 0.69), indicating comparable cyst burdens in the brain. Mice were treated for 2 weeks, starting at 14 wpi when the parasite burden is expected to peak ^33^.

We investigated the impact of blocking the Tim-3 pathway on body weight. Mice were weighed every other day during the treatment period and for up to two weeks afterward. The weight data were analyzed using repeated measures ANOVA with group as a between-subject factor and time as a within-subjects factor. The results showed a significant group effect (F (1, 26) = 15.13, p = 0.0046) and an interaction between time and group (F (13, 104) = 2.66, p = 0.0029). As shown in Fig. 2, isotype control mice gained weight over time, whereas Tim-3-treated mice maintained or slightly lost weight.

**Figure 2.**
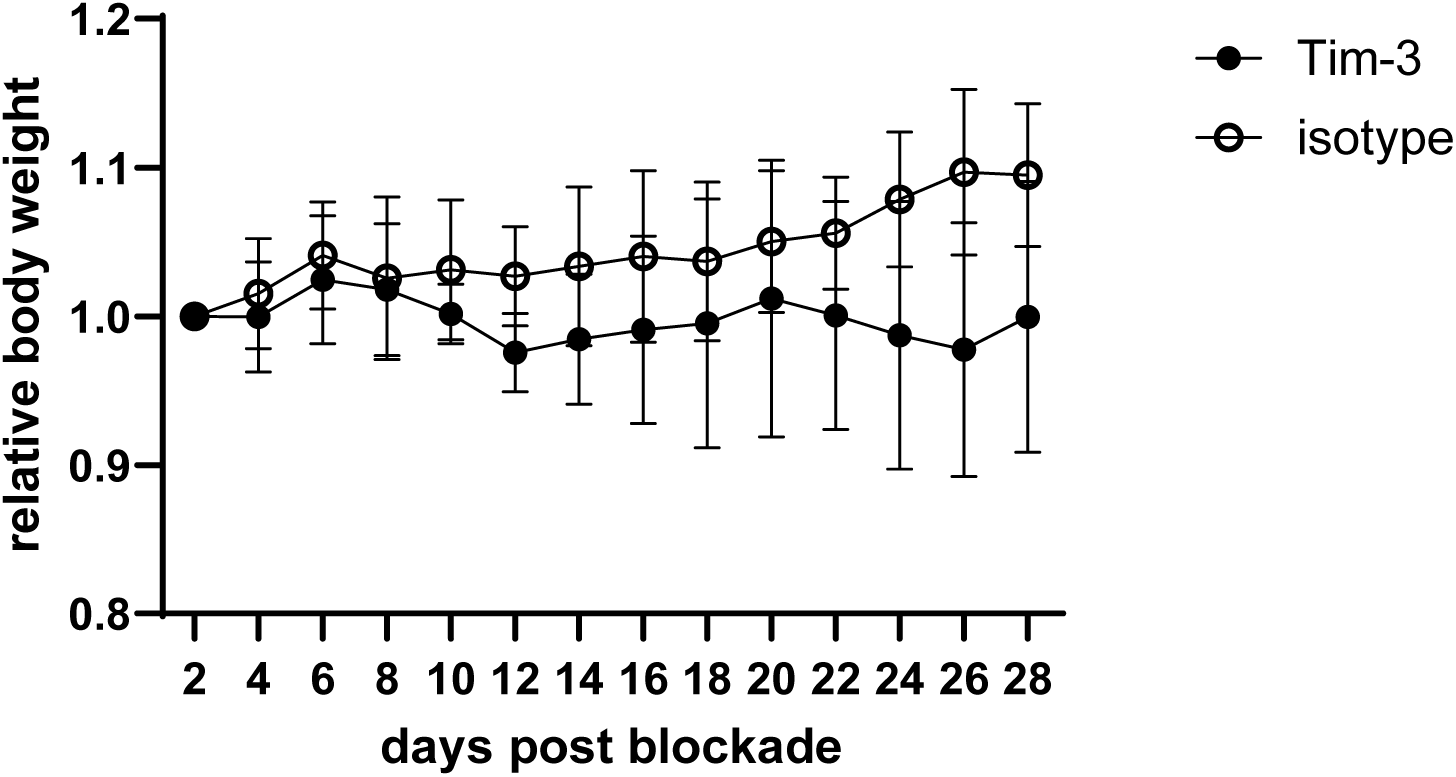
Tim-3-treated mice had poor weight gain. Mice were weighed every other day for four weeks after blocking Tim-3 signaling, and the weights were analyzed using repeated measures ANOVA. The blockade significantly affected body weight and showed an interaction with time. All the body weights were expressed as percentage change relative to pre-block weights. Data are shown as mean ± SD.

Mice were sacrificed 2 weeks after the final injection of αTim-3 antibody, and the number of tissue cysts in the brain was counted. As shown in Fig. 3A, the isotype control group (IgG2b) had a mean of 201 cysts per brain, while the αTim-3-treated group had 75 cysts, representing a 62% reduction (P < 0.0001). The average size of the cysts was smaller in the αTim-3 group (16 ± 15.2 μm) compared to the isotype control (27 ± 5.7 μm); however, this difference was not statistically significant (p = 0.134, Fig. 3B) due to considerable variation. We also used qPCR to assess the cyst burden by measuring the expression of the major bradyzoite antigen, BAG1, in brain homogenates. Due to our low-cyst-burden model, the accuracy of absolute quantification is susceptible to substantial sampling error. Therefore, we performed BAG1 qPCR three times on brain homogenates for each mouse as described in the Methods section. The difference between groups was analyzed by the Mann-Whitney U test (p = 0.12). The test was validated by examining the correlation between cyst number and qPCR results (r = 0.63, p = 0.0452). We also measured the tachyzoite-specific gene SAG1 and found it undetectable in both groups, indicating no interconversion from bradyzoites to tachyzoites.

**Figure 3.**
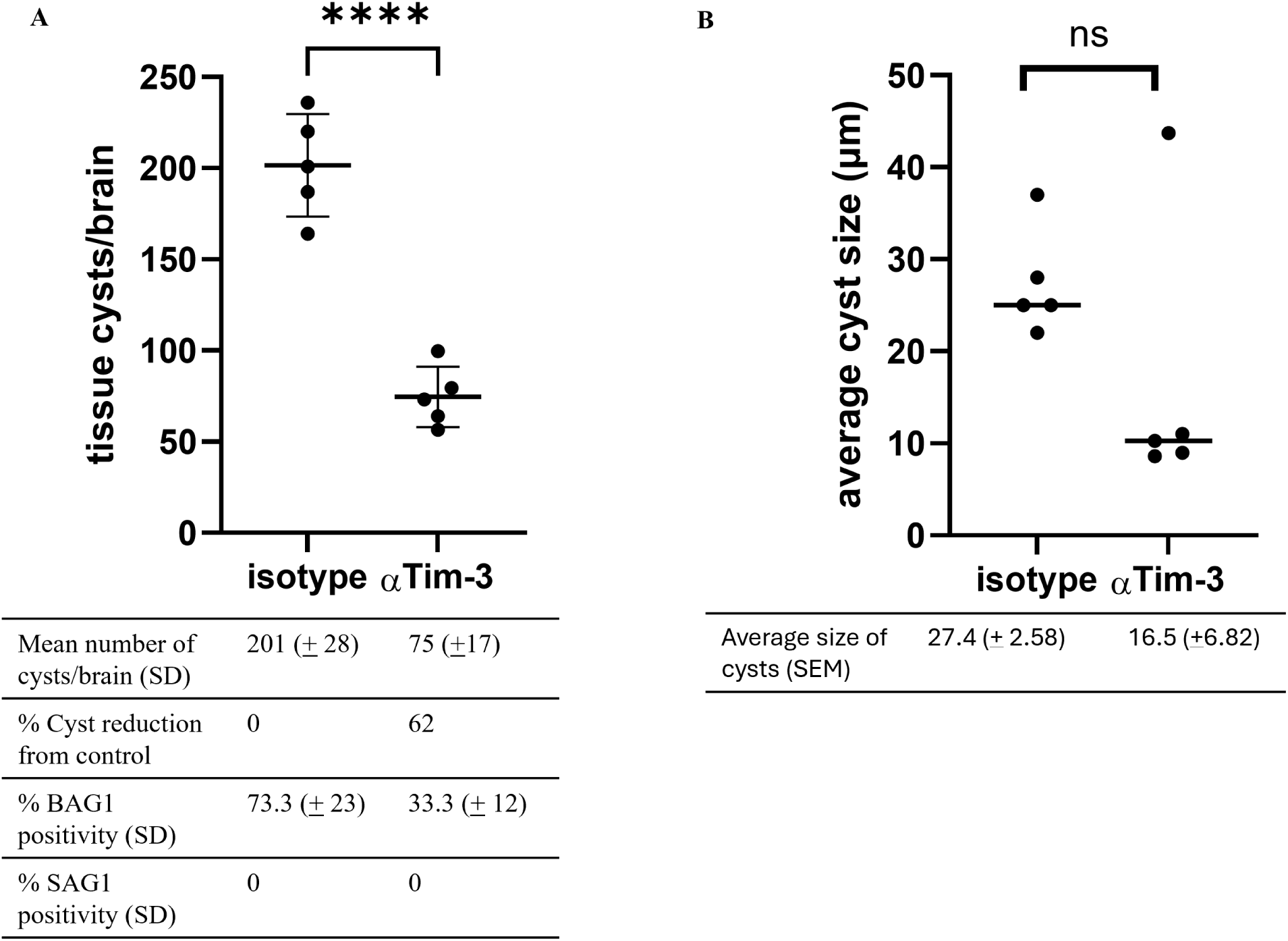
Efficacy of αTim-3 mAb against latent *T. gondii* infection. CD-1 mice were infected intraperitoneally with 500 tachyzoites of the GT1 strain and treated with αTim-3 mAb or isotype control for 2 weeks at 14 wpi. Two weeks after treatment, the mice were sacrificed, and brain tissue cysts were measured. **(A)** αTim-3 mAb treatment reduced the number of *T. gondii* cysts by 62% compared to the isotype control. **(B)** The mean size of cysts in the Tim-3 group appeared smaller, but the differences were not statistically significant. P values were calculated using Student’s t-test (A) and Mann Whitney test (B). Data are shown as mean ± SD, with individual data points. ns, p > 0.05; ****p < 0.0001.

### Blockade of Tim-3 increased the serum levels of several cytokines

To assess the effect of the Tim-3 blockade on cytokine production, we measured 23 cytokines and chemokines in serum using a Bio-Plex multiplex assay (Table 1). Compared to the isotype control, Tim-3 blockade resulted in significant increases in several cytokines, including IL-12p70 (p = 0.029), IFN-γ (p = 0.048), IL-2 (p = 0.050), CXCL1 (p = 0.045), and IL-9 (p = 0.040) and a trend for increases in Eotaxin (p = 0.061), IL-13 (p = 0.067), TNF-α (p = 0.070), and MCP-1 (p = 0.090). Notably, IL-13 showed the highest increase at 8-fold, followed by IFN-γ and Eotaxin, which increased by 6.1-fold and 6-fold, respectively. IL-9 also showed a large fold change (20-fold), but results are skewed due to undetectable levels in some mice. Furthermore, higher levels of IL-13 (r = −0.96, p=0.003), IL-2 (r = −0.86, p=0.024), and CXCL1 (r = −0.96, p=0.003) correlated with a lower burden of brain cysts. No significant changes were observed for the remaining 14 cytokines in the multiplex panel.

**Table 1.**
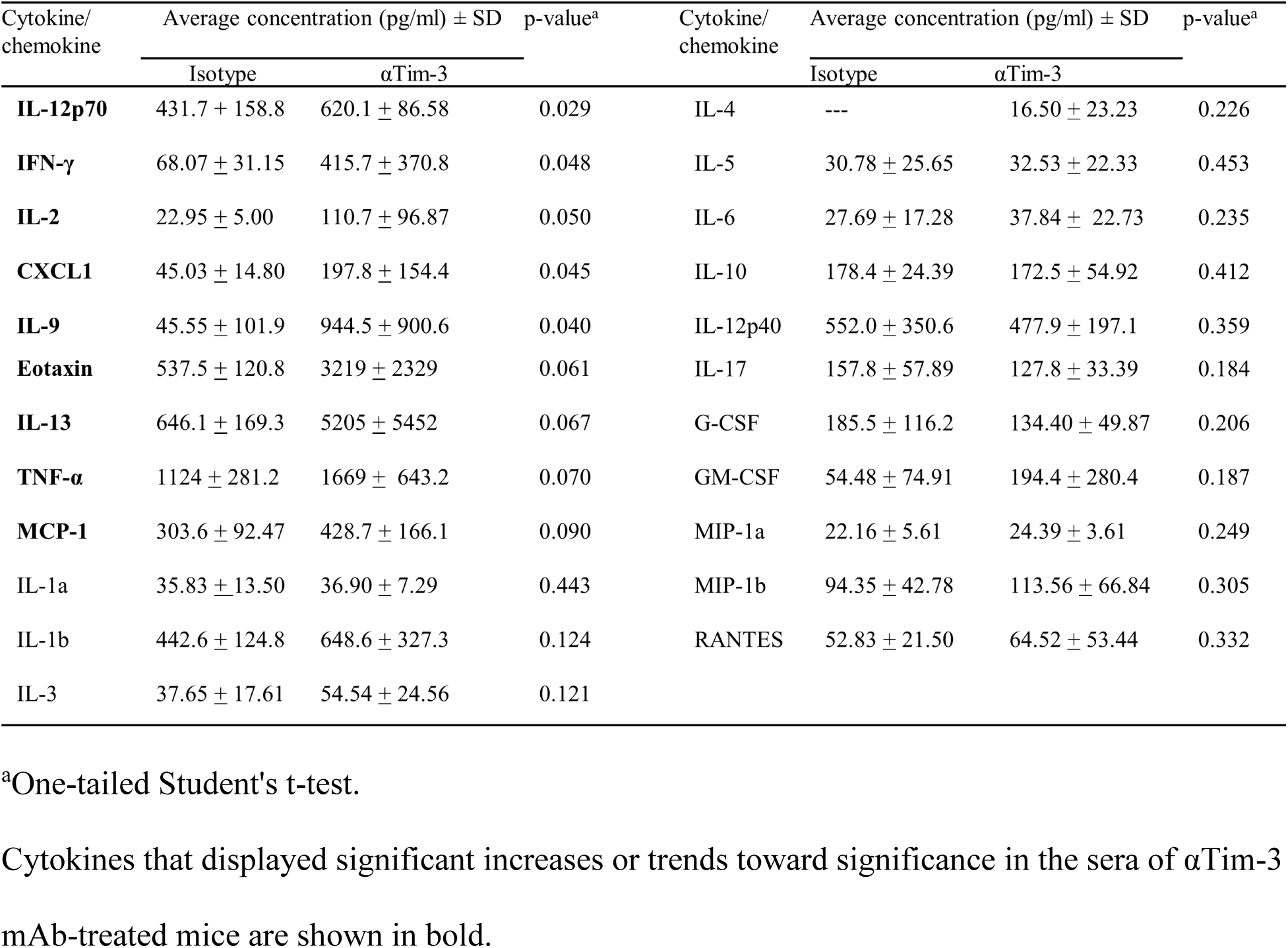
The serum concentrations of 23 cytokines in *T. gondii*-infected mice between αTim-3 mAb and isotype control treatments.

### Blockade of Tim-3 increased leukocyte infiltration into brain ventricles

In our previous study of PD-L1 blockade, we found that eliminating tissue cysts is linked to increased immune cell infiltration into the brain through cerebrospinal fluid (CSF)-filled compartments ^20^. To investigate whether a similar process occurs when blocking Tim-3, we conducted an immunohistochemical examination of immune cells in the brain. Using antibodies against CD3 (T cells) and CD14 (monocytes/macrophages), we found that mice treated with αTim-3 had increased numbers of CD3+ and CD14+ cells compared to those treated with isotype controls (Fig. 4A-B, and Supplementary Fig. S1). These immune cells were primarily located within the choroid plexus, subependymal tissue, and proximal brain parenchyma. No immune cells were observed in brain regions distant from the ventricles. In contrast, brain sections from isotype-treated mice displayed only a few cells in the ventricles.

**Figure 4.**
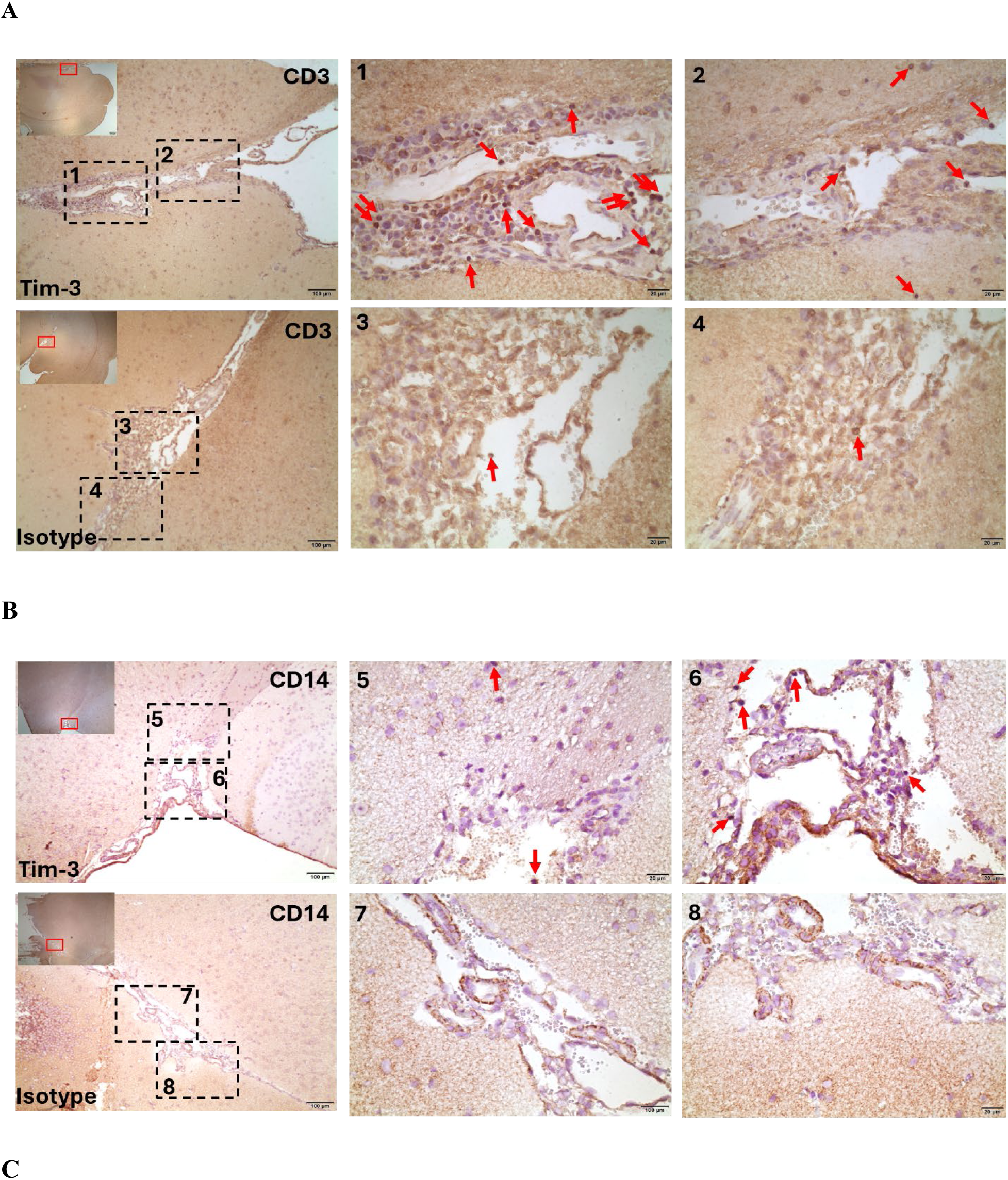
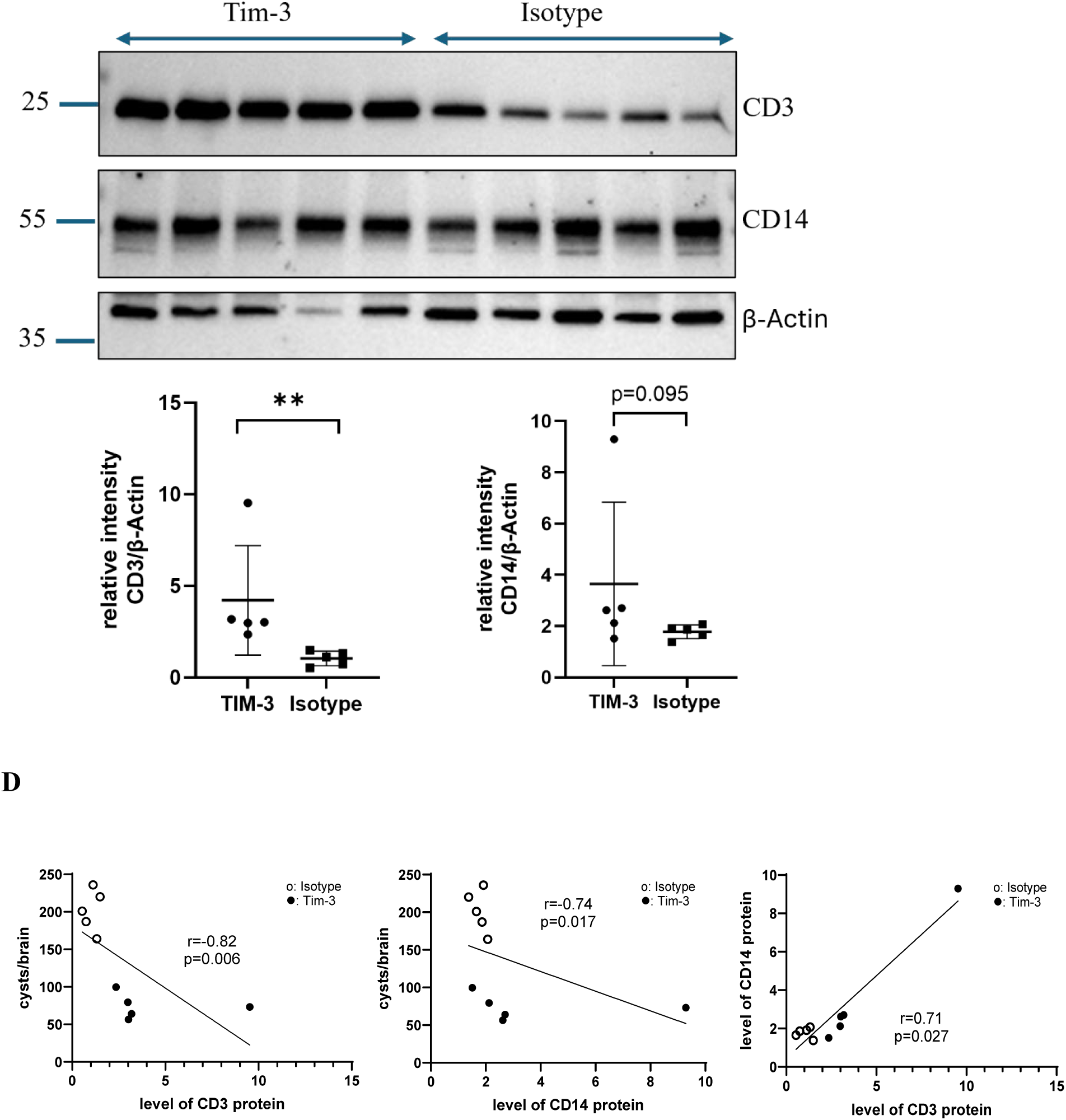
Blocking Tim-3 signaling led to increased leukocyte infiltration into the brain, which was correlated with reduced parasite burden. Compared to isotype control, there was an increased accumulation of **(A)** CD3+ and **(B)** CD14+ cells in the choroid plexus, subependymal tissue, and proximal brain parenchyma of Tim-3-treated mice. First panels: gross anatomical areas of interest (insert red box at 4× magnification) and areas marked with dotted line box (10× magnification). Leukocytes are identified at higher magnification (red arrows) in the second and third panels (40× magnification). Scale bar =100 μm (first panels), and Scale bar = 20 μm (second and third panels). The staining was repeated in two additional mice, and similar results were observed. **(C)** Western blot analysis compared levels of CD3 and CD14 protein in brain homogenates of mice treated with αTim-3 (n = 5) and isotype control (n = 5), with β-actin as a loading control. Histograms indicate densitometric analysis of blots, expressed as means ± SD, with individual data points. Significance was determined with Mann Whitney test (**p < 0.01). **(D)** Spearman’s correlation analysis between levels of CD3 and CD14 protein and the number of brain cysts in both the Tim-3 and isotype groups, as well as the correlation between the levels of CD3 and CD14 protein (n=10). Black dot: αTim-3 mice; circle: isotype mice.

We then quantified the expression of CD3 and CD14 proteins in brain homogenates from mice in both groups. Western blot analysis showed that CD3 expression was significantly higher in the αTim-3-treated mice compared to the isotype-treated group (mean: 4.215 vs. 1.043, p = 0.0464, Fig. 4C). However, there was no significant difference in CD14 expression (p > 0.5; Fig. 4C).

We also observed negative correlations between the number of brain cysts and the levels of CD3 and CD14 proteins across both Tim-3 and isotype-treated mice (CD3: r = -0.82, p = 0.006; CD14: r = -0.74, p = 0.017, Fig. 4C). Additionally, there was a positive correlation between CD3 and CD14 levels (r = 0.71, p = 0.027, Fig. 4C).

### Blockade of Tim-3 led to the restoration of P2RY12 expression

Research has shown that the extent of brain inflammation is correlated with parasite burden ^47^. Given that the Tim-3-treated mice exhibited a reduced parasite burden, we investigated whether this would lead to a decrease in immune-related responses, such as microglial activation.

Microglia activation is a hallmark of chronic *T. gondii* infection ^47^. We identified microglia using the markers IBA1 and P2RY12. IBA1 is a mark of both resident microglia and infiltrating macrophages and is upregulated upon activation, whereas P2RY12 is specific for homeostatic microglia ^15^.

We performed double staining with antibodies against P2RY12 and IBA1 on brain sections, focusing on the cortical region (Fig. 5A and Supplementary Fig. S2). P2RY12 protein primarily localized in the cytoplasm and ramified processes. Tim-3-treated mice exhibited more intense P2RY12 staining than isotype-treated mice, with an MFI of 5.49 (± 1.87) versus 2.02 (± 0.46) and a percentage area of 2.15 (± 0.73) compared to 0.79 (± 0.18) (p = 0.005). In contrast, Tim-3-treated mice had reduced IBA1 staining, with an MFI of 3.17 (± 0.72) versus 4.35 (± 0.64) and a percentage area of 1.24 (± 0.28) compared to 1.71 (± 0.25) (p = 0.013). Colocalization analyses for P2RY12 and IBA1 revealed Pearson’s coefficient of 56.6.7% (± 0.03) in the Tim-3 mice, which decreased compared to 67.6% (± 0.11) in the isotype control (p = 0.050). In support of this, microglia expressing solely P2RY12, without IBA1, were present in Tim-3 mice (Fig. 5A and Supplementary Fig. S2). We also performed further analysis using western blot to quantify the expression of P2RY12. As shown in Fig. 5B, the anti-P2RY12 antibody detected two polypeptide bands in brain homogenates: one band at approximately 58 kDa, which is considered to represent full-length P2RY12, and another band at approximately 28 kDa, which is presumed to be a cleavage fragment of the full-length P2RY12 polypeptide ^48^. Semi-quantitative measurements of band intensities, normalized against β-actin levels, indicated a trend towards increased levels of both the 58 kDa and 28 kDa polypeptides in the Tim-3 group compared to the isotype control group. The mean for the 58 kDa polypeptide was 0.343 (Tim-3) versus 0.118 (control, p = 0.054), and for the 28 kDa polypeptide, it was 3.564 (Tim-3) versus 1.320 (control, p = 0.068; see Fig. 5B). However, there was no significant difference in the levels of the IBA1 protein between the two groups (p = 0.171, Fig. 5B).

**Figure 5.**
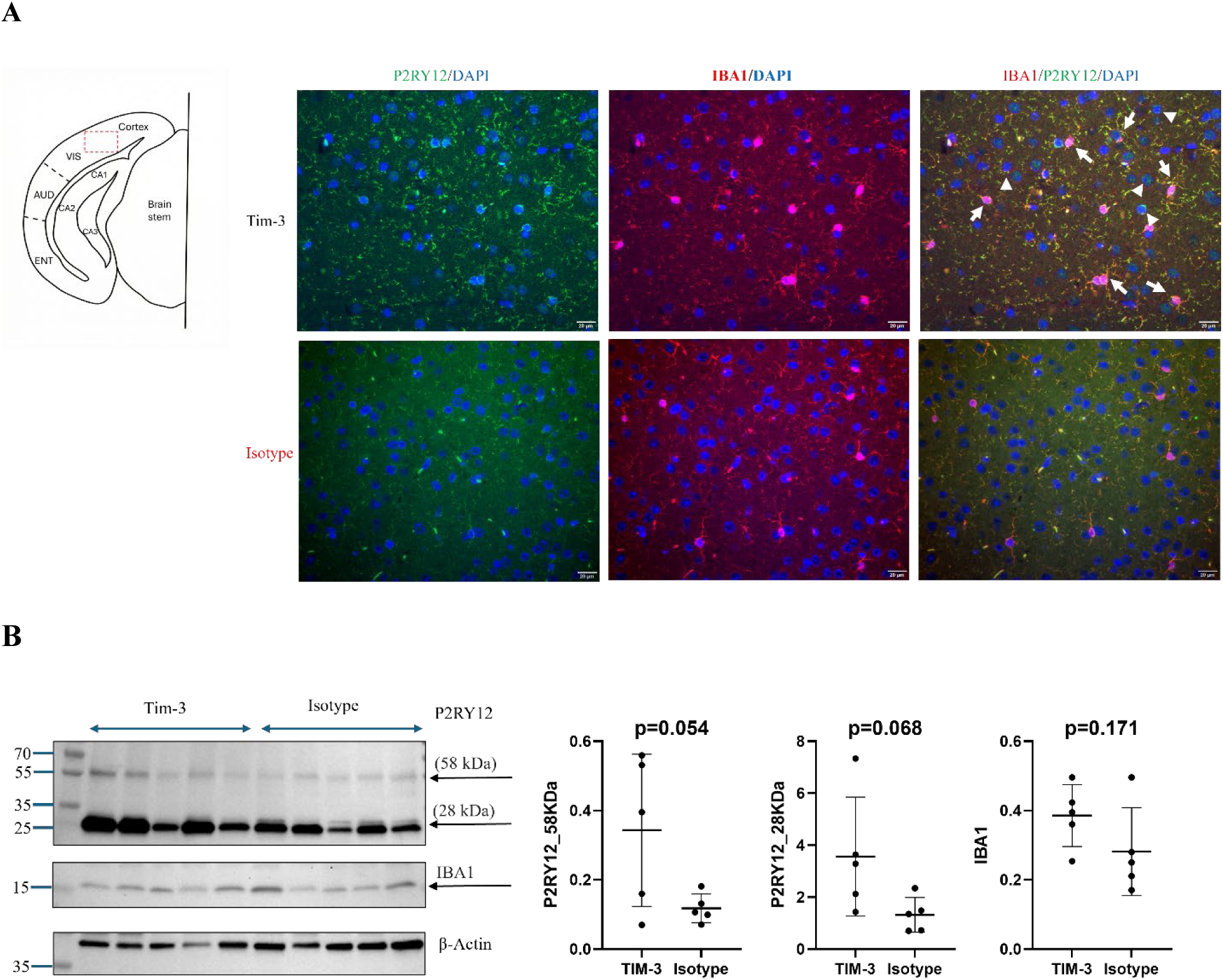
Blocking Tim-3 signaling restored the expression of P2RY12 protein. **(A)** An anatomical diagram shows section location. VIS: Visual area; AUD: Auditory area; ENT: Enthorhinol area; CA1: Cornu ammonis area 1; CA2: Cornu ammonis area 2; CA3: Cornu ammonis area 3; DG: Dentate Gyrus. Coronal sections of the mouse brain from the Tim-3 or isotype group underwent dual-immunofluorescence staining for P2RY12 and IBA1 at the visual cortex. The P2RY12 staining was localized at the cell body as well as along cell processes. In Tim-3-treated mice, P2RY12 staining was observed in both IBA1+ microglial cells (white arrows) and those lacking IBA1 (arrowhead). Scale bar = 20 μm for all panels. **(B)** Western blot analysis compared P2RY12 polypeptide and IBA1 protein levels in brain homogenates of mice treated with αTim-3 (n = 5) and isotype control (n = 5), with β-actin as a loading control. Histograms indicate densitometric analysis of blots, expressed as means + SD, with individual data points. Significance was determined with a Student’s t-test.

### Blockade of Tim-3 diminished the expression of Tim-3

Since Tim-3 expression correlates with parasite burden, we investigated whether its expression in Tim-3-treated mice decreased in correlation with reduced parasite burden. Our results revealed a significant decrease in Tim-3 mRNA expression in the brain homogenates of Tim-3-treated mice when compared to the isotype group (Fig. 6A). We then visualized Tim-3 expression in the brain by performing double immunofluorescence analysis with antibodies against Tim-3 and the microglia markers, as microglia are the primary producers of Tim-3 ^49^. We used microglial inflammatory/homeostatic markers (Tim-3/P2RY12). Tim-3 was found to be expressed on the membrane, exhibiting a dot-like pattern. Mice treated with Tim-3 showed decreased staining for Tim-3 compared to isotype-treated mice, with an MFI of 1.09 (± 0.40) versus 1.60 (± 0.57) and a percentage area of 0.43 (± 0.16) versus 0.63 (± 0.22), Ps = 0.106; Fig. 6B-C). There was an inverse relationship between Tim-3 and P2RY12 expression; higher P2RY12 levels were linked to lower Tim-3 levels in Tim-3-treated mice, and vice versa in isotype-treated mice (Fig. 6B and Supplementary Fig. S3A). In contrast, an opposite pattern was found between Tim-3 and IBA1 expression. elevated Tim-3 expression was associated with higher IBA1 staining and more extensive branching in isotype-treated mice, while lower IBA1 levels correlated with reduced Tim-3 staining in Tim-3-treated mice (Fig. 6C and Supplementary Fig. S3B).

**Figure 6.**
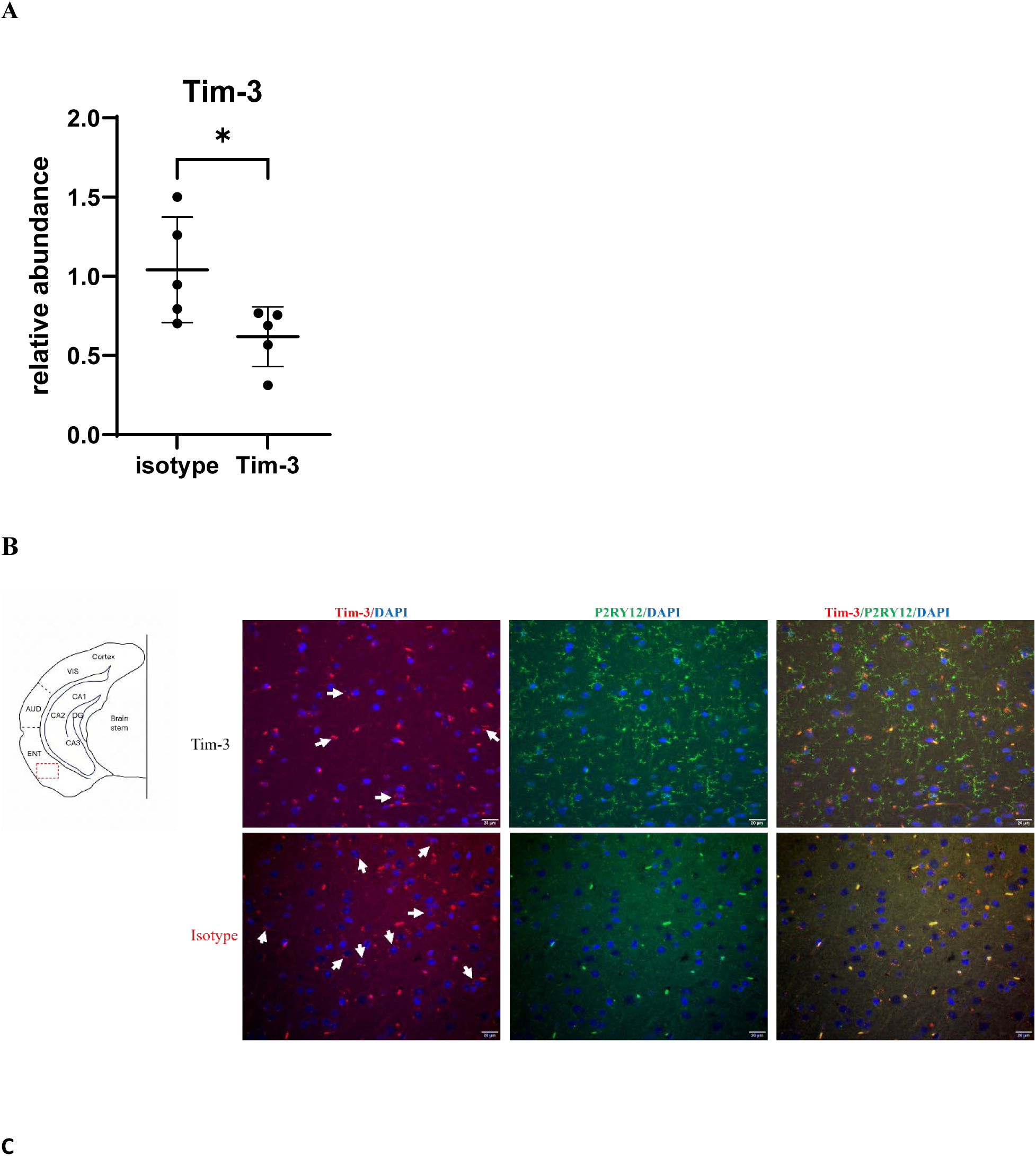
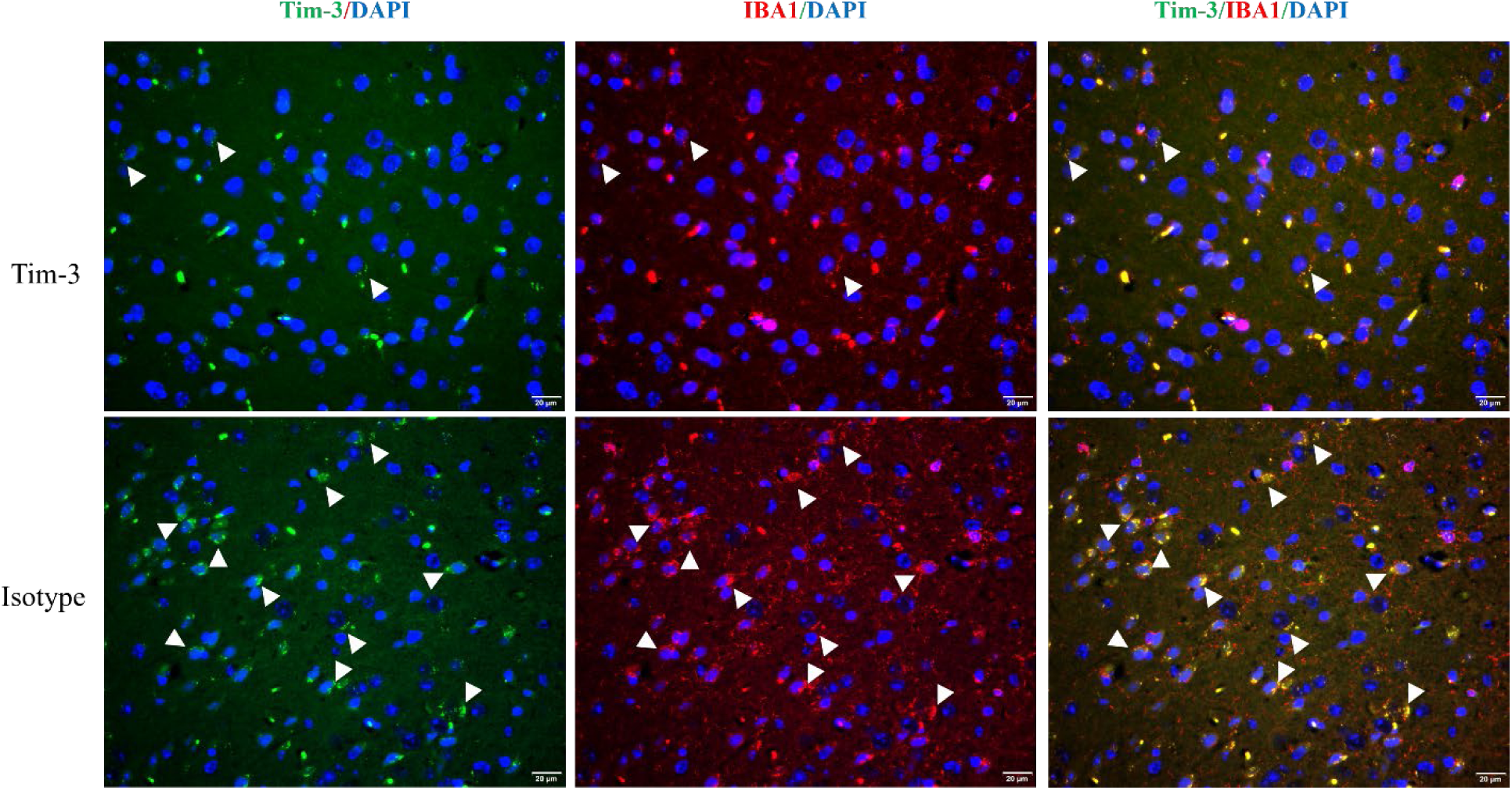
Blocking Tim-3 signaling led to reduced Tim-3 expression (A), and Tim-3 expression was negatively correlated with P2RY12 (B) and positively correlated with IBA1 expression (C). **(A)** mRNA expression levels of Tim-3 measured by RT-qPCR in brain homogenates from mice treated with Tim-3 and isotype mAb. P values were calculated using Student’s t-test (*p < 0.05). Data are shown as mean ± SD, with individual data points. **(B)** On the left side, an anatomical diagram illustrates the locations for sections (B) and (C). VIS: Visual area; AUD: Auditory area; ENT: Enthorhinol area; CA1: Cornu ammonis area 1; CA2: Cornu ammonis area 2; CA3: Cornu ammonis area 3; DG: Dentate Gyrus. Coronal sections of the mouse brain were subjected to dual immunofluorescence staining of TIM-3 with P2RY12 at the entorhinal cortex. The staining of Tim-3 was negatively associated with P2RY12-positive cells. The arrows indicate the cells expressing Tim-3. **(C)** Coronal sections of the mouse brain were subjected to dual immunofluorescence staining of TIM-3 with IBA1. The staining of Tim-3 was positively associated with IBA1-positive cells. The arrowheads indicate the cells expressing Tim-3, IBA1, and both. Scale bar = 20 μm for all panels.

## Discussion

Tissue cysts, characteristic of chronic *T. gondii* infection, are mainly found in the brain and are resistant to currently available treatments. Previous studies have highlighted the significance of T-cell exhaustion and the potential of checkpoint inhibitors in treating chronic infections such as *T. gondii* ^17^. Using a mouse model of chronic *T. gondii* infection, we investigated the effects of blocking the immune checkpoint, Tim-3, a specific marker for Th1 responses. There was an increased expression of Tim-3 in both cerebral and peripheral tissues, and this expression was correlated with MAG1 antibody levels, a surrogate marker for the brain parasite burden ^33^.

Inhibiting Tim-3 signaling resulted in reactivation of the host’s immune system, a 62% decrease in tissue cysts, and a reduction in brain inflammation. Our study provides proof of concept for blocking Tim-3 signaling as a therapeutic approach for chronic toxoplasmosis.

Our results indicate that blocking Tim-3 signaling reactivated the host’s immune system, as evidenced by increased serum cytokine levels, heightened brain leukocyte infiltration, and reduced Tim-3 expression. These findings are consistent with our previous study on blocking PD-L1 in *T. gondii*-infected mice ^20^. Monoclonal antibodies generally cannot cross the blood-brain barrier (BBB), which raises concerns about their effectiveness in treating tissue cysts in the brain. However, our research suggests that crossing the BBB is not necessary for their effectiveness. Instead, these antibodies enhance overall immunity, which facilitates the trafficking of immune cells, such as CD3+ and CD14+ cells, through the ventricles to the brain. Since the infiltration of immune cells from the periphery is crucial for combating chronic *T. gondii* infections ^50^, this increase likely contributes to the clearance of tissue cysts. Indeed, higher levels of immune cell infiltration were correlated with lower numbers of tissue cysts in the brain. However, we recognize the limitations in quantifying increases in CD3⁺ and CD14⁺ cells, as immunofluorescence images are qualitative, and Western blots may reflect changes in cell number or protein expression levels. Anti-Tim-3 mAb-treated mice maintained or slightly lost weight during treatment, but no additional toxicities or side effects were observed. Tim-3 blockade, either alone or in combination with CTLA-4 blockade, has been associated with reduced body weight growth in a mouse pregnancy model but not weight loss ^51,52^.

A higher antigen load likely leads to the continuous stimulation of parasite-specific T cells, resulting in an exhausted phenotype with the expression of inhibitory molecules ^17,53^. Our study found a positive correlation between parasite burden and Tim-3 expression in the prefrontal cortex, striatum, heart, and liver of *T. gondii*-infected mice. This finding aligns with previous research, which has shown that Tim-3 levels are associated with viral load and disease progression ^54–56^. Notably, Tim-3 expression decreased in the brains of mice treated with anti-αTim-3 mAb, likely related to a lower parasite burden. However, our study did not identify the phenotype of immune cells with decreased Tim-3 expression. Tim-3 is an inducible marker whose expression is downregulated on human NK cells in response to cancer targets ^57^. In a mouse model of head and neck squamous cell carcinoma (HNSCC), the percentage of Tim-3+ cells decreased in both the draining lymph nodes and the spleen following Tim-3 blockade ^58^. Additionally, a study found that patients infected with Plasmodium vivax exhibited a lower frequency of Tim-3–expressing CD4+ and CD8+ T cells after successful antimalarial treatment compared to before the treatment ^59^.

Previous research indicates that blocking Tim-3 signaling enhances cytokine production by lymphocytes during chronic infections such as HIV and Plasmodium ^27,54,59^. Elevated levels of cytokines, including IFN-γ, IL-12p70, IL-2, CXCL1, IL-9, TNFα, MCP-1, eotaxin and IL-13, were observed in Tim-3-treated mice. The increases in IFN-γ, IL-2, IL-12p70, TNFα, and MCP-1 levels are consistent with an enhanced Th1-mediated immunity and characteristic of the response to *T. gondii* infection ^60^. Notably, IFN-γ increased by over 6-fold, which is crucial for combating chronic *T. gondii* infection. IL-9 is a pleiotropic cytokine primarily produced by Th9 cells. IL-9 stimulates mast cell proliferation and activation, enhances survival and activation of T cells, and promotes protection at mucosal surfaces. Although known to play an important role in helminth infections, IL-9 has not previously been implicated in toxoplasmosis. While pro-inflammatory cytokines increased, there was also an 8-fold increase in the anti-inflammatory cytokine IL-13. Research has shown that IL-13 production is elevated in mast cells by anti-Tim-3 antibodies after antigen-dependent activation and IgE sensitization ^61,62^. Increased levels of IL-13 can shift microglia from a pro-inflammatory M1-like phenotype to an anti-inflammatory M2a phenotype, promoting repair, regeneration, and tissue remodeling ^14^. In a rat model of intracerebral hemorrhage, the downregulation of Tim-3 promoted the conversion of M1 microglia into M2 microglia ^63^. A rise in anti-inflammatory molecules may limit immune-related pathology ^19^. Eotaxin (CCL11) is a chemokine that recruits eosinophils. However, its receptor CCR3 is expressed on a wide range of immune cells including T cells and microglia, and it may play a role in neuroinflammation ^64,65^. CXCL1 is a chemoattractant for neutrophils. A phenotypically and transcriptionally unique population of neutrophils in the CNS, associated with neuronal protection, has been reported in a murine model of chronic *T. gondii* infection ^66^. A limitation of our study is that systemic measurement of cytokines may not reflect CNS levels.

Mice treated with anti-αTim-3 mAb showed a trend toward increased expression of the P2RY12 protein in microglia. P2RY12 is involved in microglial motility and migration, and its expression is used as a marker for non-activated or homeostatic microglia. A decrease in P2RY12 expression is associated with activated microglia ^15^. One study reported that P2RY12 expression is restored as the distance from amyloid plaque increases. Microglia that are in direct contact with mature plaques do not express P2RY12, while those nearby show low or no expression, and microglia farther away express it fully ^48^. In human multiple sclerosis (MS), microglia lose P2RY12 expression and adopt a pro-inflammatory phenotype ^13^. Therefore, the trend toward increased expression of P2RY12 in Tim-3 treated mice would be consistent with a reduction in inflammation. This is supported by an increase in anti-inflammatory cytokines, such as IL-13, and a decrease in Tim-3 expression in these mice. While a significant decrease in IBA1 immunostaining was noted in Tim-3-treated mice, the protein levels in brain homogenates were comparable to those in isotype controls. IBA1 serves as a marker for microglia and macrophages, which does not allow for exclusive identification of microglia. The increased presence of CD14+ macrophages in the Tim-3 group complicates this distinction. Thus, the lack of difference in IBA1 levels may not accurately reflect the actual situation and require further investigation.

Currently, the therapeutic potential of targeting Tim-3 is being tested in cancer, where its co-blockade with other checkpoint receptors is being investigated in the clinic, with promising results in patients with anti-PD1-refractory disease ^24^. Our previous research has demonstrated that blocking the PD-L1 pathway significantly reduces the number of tissue cysts ^20,21^. Tim-3 and PD-1 are potent immunoinhibitory molecules involved in immune tolerance, autoimmune responses, and the evasion of immune responses. Our findings suggest that immune checkpoint inhibition could be an effective treatment for chronic toxoplasmosis. As immunosuppression for solid organ and stem cell transplant patients becomes more frequent, the incidence of *T. gondii*-related diseases is likely to increase. There is an urgent need to clear tissue cysts to avoid severe complications, such as encephalitis and pneumonitis, in patients with weakened immune systems. Insights gained from this research may be relevant to other brain-infecting pathogens, considering the limitations of neurotropic mouse models.

## Acknowledgments

The authors thank Drs. Robert Yolken for critical reading of the manuscript and Junying Zheng for technical support.

## Authors’ contributions

JX and RPV designed the experiments. YL and JH performed experiments. JX, RPV, and ES analyzed the data. JX, RPV, and ES wrote the paper.

## Data availability

Any additional information is available from the lead contact on request.

