## supplemental figures S1-S3 for "Blockade of Tim-3 pathway in a mouse model of Toxoplasmosis: impact on brain leukocyte infiltration, parasite burden, and neuroinflammation"

Figure legends

**Figure. S1. Blocking Tim-3 signaling led to increased leukocyte infiltration into the brain.** Compared to isotype control, there was an increased accumulation of **(A)** CD3+ and **(B)** CD14+ cells in the choroid plexus, subependymal tissue, and proximal brain parenchyma of Tim-3-treated mice. First panels: gross anatomical areas of interest (insert red box at 4× magnification) and areas marked with red box (10× magnification). Leukocytes are identified at higher magnification (red arrows) in the second and third panels (40× magnification). Scale bar =100 μm (first panels), and Scale bar = 20 μm (second and third panels).

**Figure. S2. Blocking Tim-3 signaling restored the expression of P2RY12 protein.** Coronal sections of the mouse brain from the Tim-3 or isotype group underwent dual-immunofluorescence staining for P2RY12 and IBA1 at the visual cortex. The P2RY12 staining was localized at the cell body as well as along cell processes. In Tim-3-treated mice, P2RY12 staining was observed in both IBA1+ microglial cells (white arrows) and those lacking IBA1 (arrowhead). Scale bar = 20 μm for all panels.

**Figure. S3. Tim-3 staining was positively correlated with P2RY12 but negatively correlated with IBA1 expression.** Coronal sections of the mouse brain were subjected to dual immunofluorescence staining of TIM-3 with P2RY12 and IBA1 at the entorhinal cortex. **(A)** The staining of Tim-3 was negatively associated with P2RY12-positive cells. The arrows indicate the cells expressing Tim-3. **(B)** The staining of Tim-3 was positively associated with IBA1-positive cells. The arrowheads indicate the cells expressing Tim-3, IBA1, and both. Scale bar = 20 μm for all panels.

Fig. S1

A

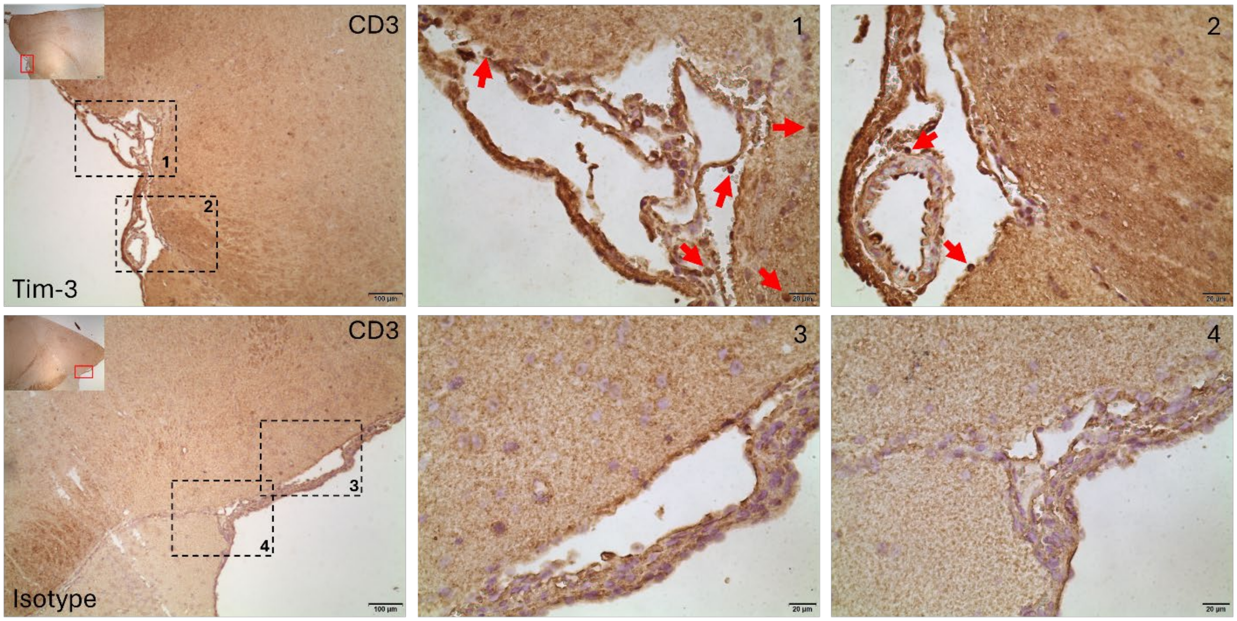


B

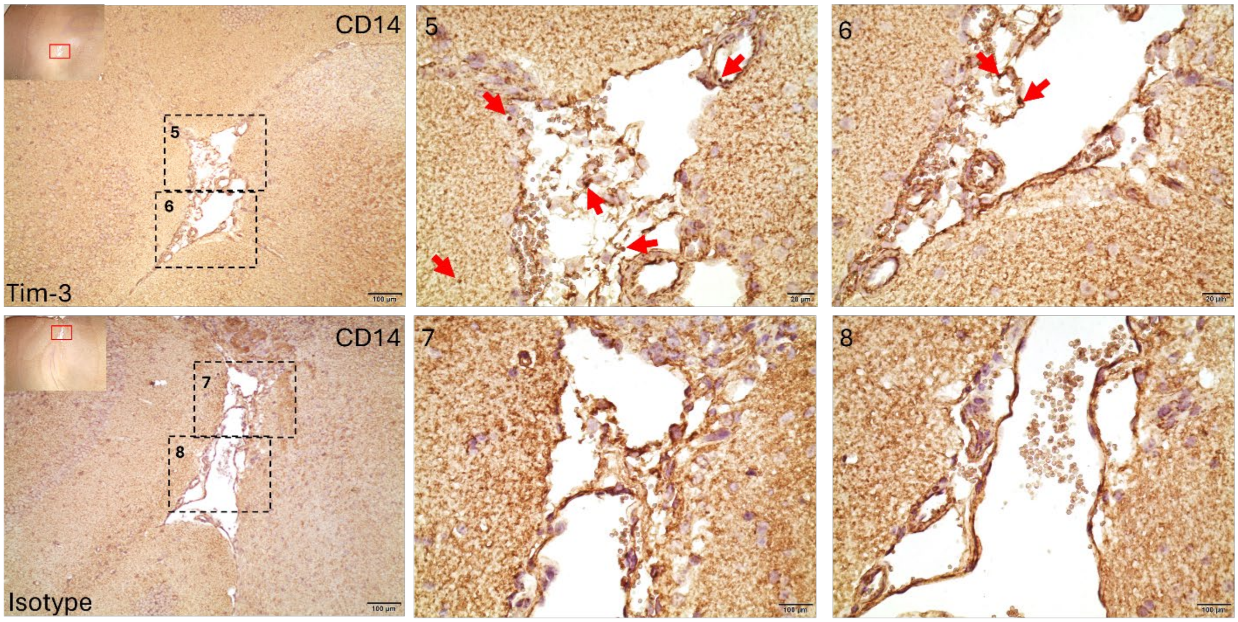

Fig. S2

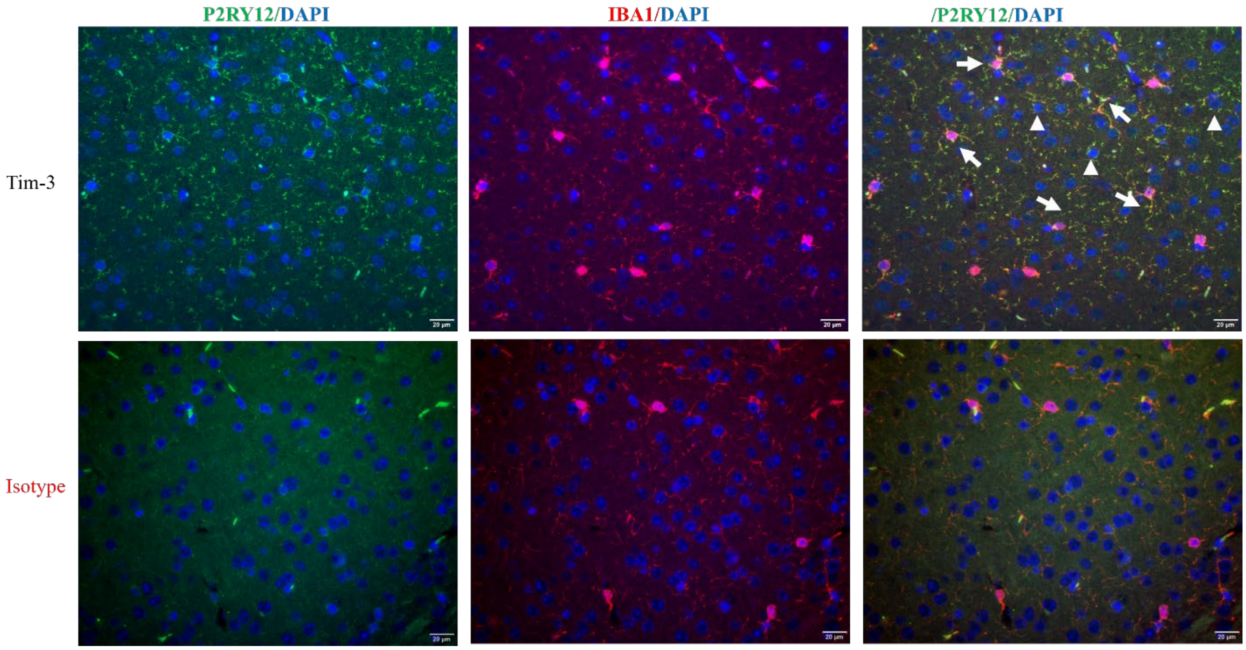

Fig. S3

A

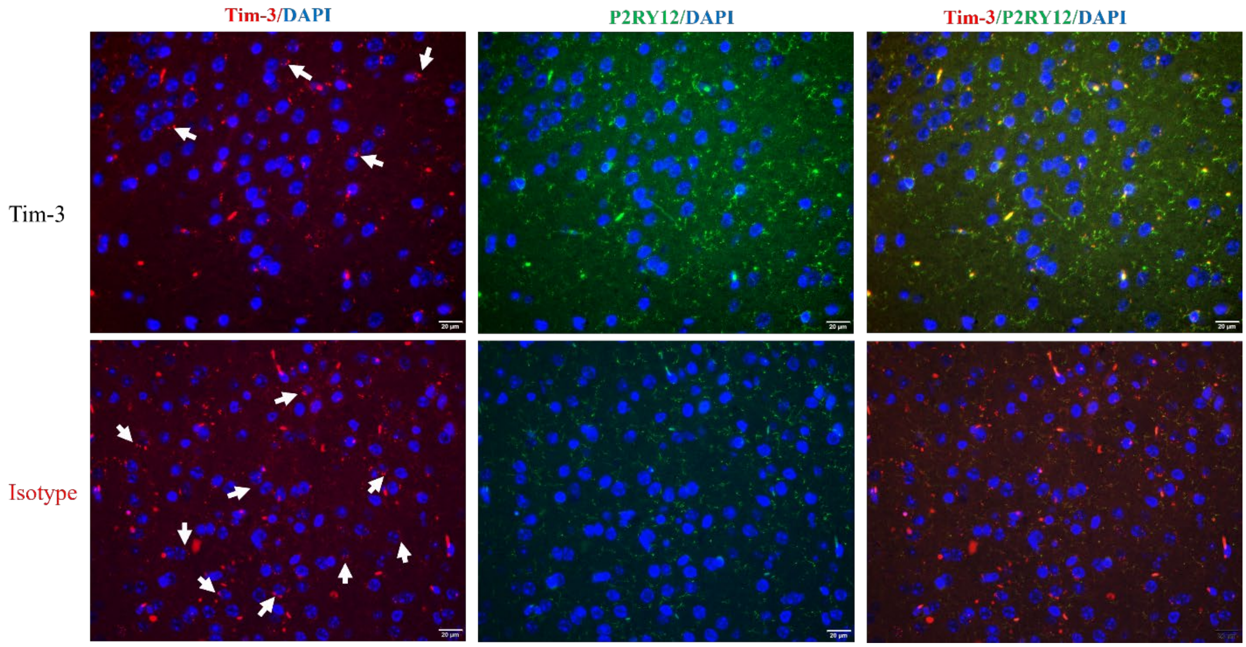
Tim-3/DAPI
P2RY12/DAPI
Tim-3/P2RY12/DAPI
Tim-3
Isotype

B

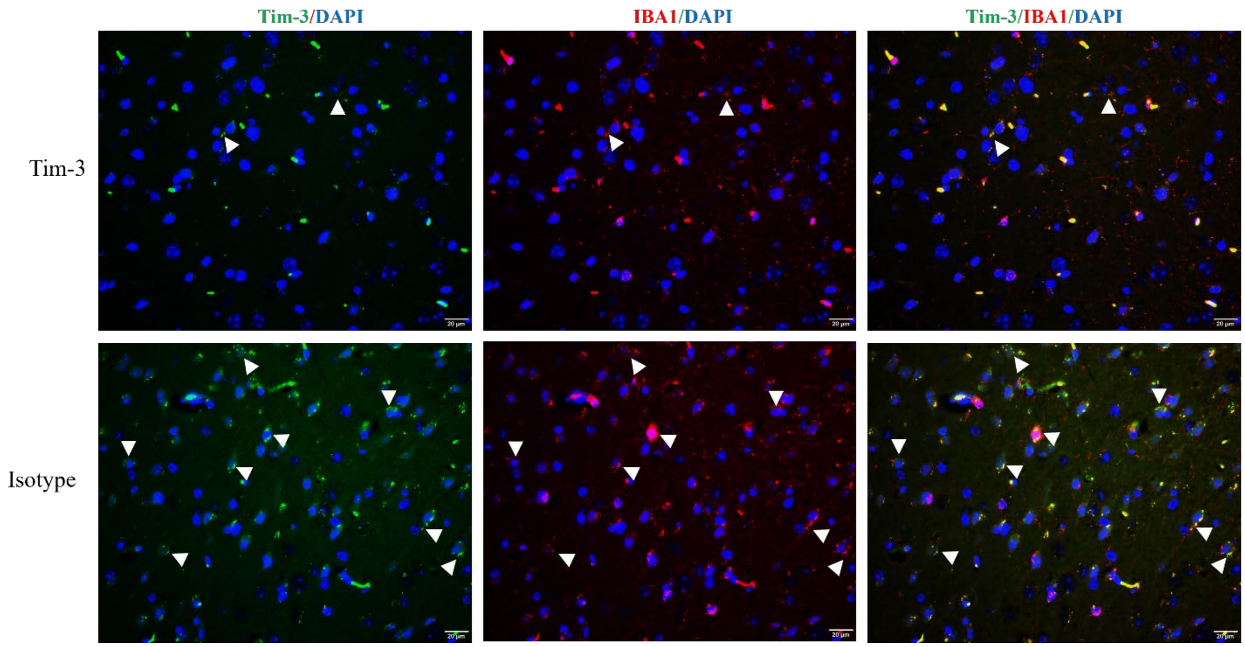
Tim-3/DAPI
IBA1/DAPI
Tim-3/IBA1/DAPI
Tim-3
Isotype
